# Recovery of deleted deep sequencing data sheds more light on the early Wuhan SARS-CoV-2 epidemic

**DOI:** 10.1101/2021.06.18.449051

**Authors:** Jesse D. Bloom

**Affiliations:** Fred Hutchinson Cancer Research Center, Howard Hughes Medical Institute, Seattle, WA, USA

## Abstract

The origin and early spread of SARS-CoV-2 remains shrouded in mystery. Here I identify a data set containing SARS-CoV-2 sequences from early in the Wuhan epidemic that has been deleted from the NIH’s Sequence Read Archive. I recover the deleted files from the Google Cloud, and reconstruct partial sequences of 13 early epidemic viruses. Phylogenetic analysis of these sequences in the context of carefully annotated existing data further supports the idea that the Huanan Seafood Market sequences are not fully representative of the viruses in Wuhan early in the epidemic. Instead, the progenitor of currently known SARS-CoV-2 sequences likely contained three mutations relative to the market viruses that made it more similar to SARS-CoV-2’s bat coronavirus relatives.

---

Understanding the spread of SARS-CoV-2 in Wuhan is crucial to tracing the origins of the virus, including identifying events that led to infection of patient zero. The first reports outside of China at the end of December 2019 emphasized the role of the Huanan Seafood Market (ProMED 2019), which was initially suggested as a site of zoonosis. However, this theory became increasingly tenuous as it was learned that many early cases had no connection to the market (Cohen 2020; Huang *et al*. 2020; Chen *et al.* 2020). Eventually, Chinese CDC Director Gao Fu dismissed the theory, stating “At first, we assumed the seafood market might have the virus, but now the market is more like a victim. The novel coronavirus had existed long before” (Global Times 2020).

Indeed, there were reports of cases that far preceded the outbreak at the Huanan Seafood Market. The *Lancet* described a confirmed case having no association with the market whose symptoms began on December 1, 2019 (Huang *et al.* 2020). The *South China Morning Post* described nine cases from November 2019 including details on patient age and sex, noting that none were confirmed to be “patient zero” (Ma 2020). Professor Yu Chuanhua of Wuhan University told the *Health Times* that records he reviewed showed two cases in mid-November, and one suspected case on September 29 (Health Times 2020). At about the same time as Professor Chuanhua’s interview, the Chinese CDC issued an order forbidding sharing of information about the COVID-19 epidemic without approval (China CDC 2020), and shortly thereafter Professor Chuanhua re-contacted the *Health Times* to say the November cases could not be confirmed (Health Times 2020). Then China’s State Council issued a much broader order requiring central approval of all publications related to COVID-19 to ensure they were coordinated “like moves in a game of chess” (Kang *et al.* 2020a). In 2021, the joint WHO-China report dismissed all reported cases prior to December 8 as not COVID-19, and revived the theory that the virus might have originated at the Huanan Seafood Market (WHO 2021).

In other outbreaks where direct identification of early cases has been stymied, it has increasingly become possible to use genomic epidemiology to infer the timing and dynamics of spread from analysis of viral sequences. For instance, analysis of SARS-CoV-2 sequences has enabled reconstruction of the initial spread of SARS-CoV-2 in North America and Europe (Bedford *et al*. 2020; Worobey *et al.* 2020; Deng *et al.* 2020; Fauver *et al.* 2020).

But in the case of Wuhan, genomic epidemiology has also proven frustratingly inconclusive. Some of the problem is simply limited data: despite the fact that Wuhan has advanced virology labs, there is only patchy sampling of SARS-CoV-2 sequences from the first months of the city’s explosive outbreak. Other than a set of multiply sequenced samples collected in late December of 2019 from a dozen patients connected to the Huanan Seafood Market (WHO 2021), just a handful of Wuhan sequences are available from before late January of 2020 (see analysis in this study below). This paucity of sequences could be due in part to an order that unauthorized Chinese labs destroy all coronavirus samples from early in the outbreak, reportedly for “laboratory biological safety” reasons (Pingui 2020).

However, the Wuhan sequences that are available have also confounded phylogenetic analyses designed to infer the “progenitor” of SARS-CoV-2, which is the sequence from which all other currently known sequences are descended (Kumar *et al*. 2021). Although there is debate about exactly how SARS-CoV-2 entered the human population, it is universally accepted that the virus’s deep ancestors are bat coronaviruses (Lytras *et al*. 2021). But the earliest known SARS-CoV-2 sequences, which are mostly derived from the Huanan Seafood Market, are notably more different from these bat coronaviruses than other sequences collected at later dates outside Wuhan. As a result, there is a direct conflict between the two major principles used to infer an outbreak’s progenitor: namely that it should be among the earliest sequences, and that it should be most closely related to deeper ancestors (Pipes *et al.* 2021).

Here I take a small step towards resolving these questions by identifying and recovering a deleted data set of partial SARS-CoV-2 sequences from outpatient samples collected early in the Wuhan epidemic. Analysis of these new sequences in conjunction with careful annotation of existing ones suggests that the early Wuhan samples that have been the focus of most studies including the joint WHO-China report (WHO 2021) are not fully representative of the viruses actually present in Wuhan at that time. These insights help reconcile phylogenetic discrepancies, and suggest two plausible progenitor sequences, one of which is identical to that inferred by Kumar *et al*. (2021) and the other of which contains the C29095T mutation. Furthermore, the approach taken here hints it may be possible to advance understanding of SARS-CoV-2’s origins or early spread even without further on-the-ground studies, such as by more deeply probing data archived by the NIH and other entities.

## Results

### Identification of a SARS-CoV-2 deep sequencing data set that has been removed from the Sequence Read Archive

During the course of my research, I read a paper by Farkas *et al*. (2020) that analyzed SARS-CoV-2 deep sequencing data from the Sequence Read Archive (SRA), which is a repository maintained by the NIH’s National Center for Biotechnology Information. The first supplementary table of Farkas *et al*. (2020) lists all SARS-CoV-2 deep sequencing data available from the SRA as of March 30, 2020.

The majority of entries in this table refer to a project (BioProject PRJNA612766) by Wuhan University that is described as nanopore sequencing of SARS-CoV-2 amplicons. The table indicates this project represents 241 of the 282 SARS-CoV-2 sequencing run accessions in the SRA as of March, 30, 2020. Because I had never encountered any other mention of this project, I performed a Google search for “PRJNA612766,” and found no search hits other than the supplementary table itself. Searching for “PRJNA612766” in the NCBI’s SRA search box returned a message of “No items found.” I then searched for individual sequencing run accessions from the project in the NCBI’s SRA search box. These searches returned messages indicating that the sequencing runs had been removed (Figure 1).

**Figure 1.**
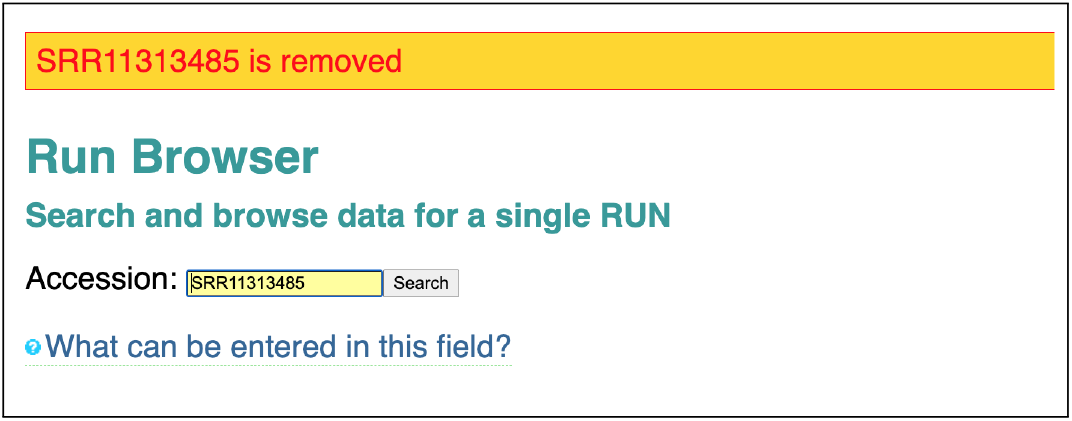
Accessions from deep sequencing project PRJNA612766 have been removed from the SRA. Shown is the result of searching for “SRR11313485” in the SRA search tool-bar. This result has been digitally archived on the Wayback Machine at https://web.archive.org/web/20210502131630/ https://trace.ncbi.nlm.nih.gov/Traces/sra/?run=SRR11313485.

The SRA is designed as a permanent archive of deep sequencing data. The SRA documentation states that after a sequencing run is uploaded, “neither its files can be replaced nor filenames can be changed,” and that data can only be deleted by e-mailing SRA staff (SRA 2021).

### The deleted data set contains sequencing of viral samples collected early in the Wuhan epidemic

The metadata in the first supplementary table of Farkas *et al*. (2020) indicates that the samples in deleted project PRNJA612766 were collected by Aisu Fu and Renmin Hospital of Wuhan University. Google searching for these terms revealed the samples were related to a study posted as a pre-print on *medRxiv* in early March of 2020 (Wang *et al.* 2020a), and subsequently published in the journal *Small* in June of 2020 (Wang *et al.* 2020b).

The study describes an approach to diagnose infection with SARS-CoV-2 and other respiratory viruses by nanopore sequencing. This approach involved reverse-transcription of total RNA from swab samples, followed by PCR with specific primers to generate amplicons covering portions of the viral genome. These amplicons were then sequenced on an Oxford Nanopore GridION, and infection was diagnosed if the sequencing yielded sufficient reads aligning to the viral genome. Importantly, the study notes that this approach yields information about the sequence of the virus as well enabling diagnosis of infection. In fact, Wang *et al*. (2020b) even list the mutations determined from this sequencing—but because this paper was published in the chemistry journal *Small*, after the sequences were removed from the SRA their existence appears to have been entirely overlooked.

The pre-print (Wang *et al.* 2020a) says the approach was applied to “45 nasopharyngeal swab samples from outpatients with suspected COVID-19 early in the epidemic.” The digital object identifier (DOI) for the pre-print indicates that it was processed by *medRxiv* on March 4, 2020, which is one day after China’s State Council ordered that all papers related to COVID-19 must be centrally approved (Kang *et al.* 2020a). The final published manuscript (Wang *et al.* 2020b) from June of 2020 updated the description from “early in the epidemic” to “early in the epidemic (January 2020).” Both the pre-print and published manuscript say that 34 of the 45 early epidemic samples were positive in the sequencing-based diagnostic approach. In addition, both state that the approach was later applied to 16 additional samples collected on February 11–12, 2020, from SARS-CoV-2 patients hospitalized at Renmin Hospital of Wuhan University.

There is complete concordance between the accessions for project PRJNA612766 in the supplementary table of Farkas *et al*. (2020) and the samples described by Wang *et al*. (2020a). There are 89 accessions corresponding to the 45 early epidemic samples, with these samples named like wells in a 96-well plate (A1, A2, etc). The number of accessions is approximately twice the number of early epidemic samples because each sample has data for two sequencing runtimes except one sample (B5) with just one runtime. There are 31 accessions corresponding to the 16 samples collected in February from Renmin Hospital patients, with these samples named R01, R02, etc. Again, all but one sample (R04) have data for two sequencing runtimes. In addition, there are 7 accessions corresponding to positive and negative controls, 2 accessions corresponding to other respiratory virus samples, and 112 samples corresponding to plasmids used for benchmarking of the approach. Together, these samples and controls account for all 241 accessions listed for PRJNA612766 in the supplementary table of Farkas *et al*. (2020).

Neither the pre-print (Wang *et al*. 2020a) nor published manuscript (Wang *et al.* 2020b) contain any correction or note that suggests a scientific reason for deleting the study’s sequencing data from the SRA. I e-mailed both corresponding authors of Wang *et al*. (2020a) to ask why they had deleted the deep sequencing data and to request details on the collection dates of the early outpatient samples, but received no reply.

### Recovery of deleted sequencing data from the Google Cloud

As indicated in Figure 1, none of the deleted sequencing runs could be accessed through the SRA’s web interface. In addition, none of the runs could be accessed using the command-line tools of the SRA Toolkit. For instance, running `fastq-dump SRR11313485` or `vdb-dump SRR11313485` returned the message “err: query unauthorized while resolving query within virtual file system module - failed to resolve accession ‘SRR11313485’”. However, the SRA has begun storing all data on the Google and Amazon clouds. While inspecting the SRA’s web interface for other sequencing accessions, I noticed that SRA files are often available from links to the cloud such as https://storage.googleapis.com/nih-sequence-read-archive/run/ <ACCESSION>/<ACCESSION>.

Based on the hypothesis that deletion of sequencing runs by the SRA might not remove files stored on the cloud, I interpolated the cloud URLs for the deleted accessions and tested if they still yielded the SRA files. This strategy was successful; for instance, as of June 3, 2021, going to https://storage.googleapis.com/nih-sequence-read-archive/run/SRR11313485/SRR11313485 downloads the SRA file for accession SRR11313485. I have archived this file on the Wayback Machine at https://web.archive.org/web/20210502130820/ https://storage.googleapis.com/nih-sequence-read-archive/run/SRR11313485/SRR11313485.

I automated this strategy to download the SRA files for 97 of the 99 sequencing runs corresponding to the 34 SARS-CoV-2 positive early epidemic samples and the 16 hospital samples from February. The SRA files for two runs (SRR11313490 and SRR11313499) were not accessible via the Google Cloud, but after I posted the first version of this manuscript as a pre-print, several individuals found archived data for these runs that had been downloaded when they were still available on the SRA, and I have used those data in the updated analysis described here (see Methods for details). I used the SRA Toolkit to get the object timestamp (`vdb-dump --obj_timestamp`) and time (`vdb-dump --info`) for all SRA files. For all files, the object timestamp is February 15, 2020, and the time is March 16, 2020. Although the SRA Toolkit does not clearly document these two properties, my *guess* is that the object timestamp may refer to when the SRA file was created from a FASTQ file uploaded to the SRA, and the time may refer to when the accession was made public.

### The data are sufficient to determine the viral sequence from the start of spike through the end of ORF10 for some samples

Wang *et al*. (2020a) sequenced PCR amplicons covering nucleotide sites 21,563 to 29,674 of the SARS-CoV-2 genome, which spans from the start of the spike gene to the end of ORF10. They also sequenced a short amplicon generated by nested PCR that covered a fragment of ORF1ab spanning sites ∼ 15,080 to 15,550. In this paper, I only analyze the region from spike through ORF10 because this is a much longer contiguous sequence and the amplicons were generated by conventional rather than nested PCR. I slightly trimmed the region of interest to 21,570 to 29,550 because many samples had poor coverage at the termini.

I aligned the recovered deep sequencing data to the SARS-CoV-2 genome using `minimap2` (Li 2018), combining accessions for the same sample, and masking regions that corresponded to the primer binding sites described in Wang *et al*. (2020a). Figure S1 shows the sequencing coverage for the 34 virus-positive early epidemic samples and the 16 hospitalized patient samples over the region of interest; a comparable plot for the whole genome is in Figure S2.

I called the consensus viral sequence for each sample at each site with coverage ≥3 and >80% of the reads concurring on the nucleotide identity. With these criteria, 13 of the early outpatient samples and 1 of the February hospitalized patient samples had sufficient coverage to call the consensus sequence at >90% of the sites in the region of interest (Table 1), and for the remainder of this paper I focus on these high-coverage samples. Table 1 also shows the mutations in each sample relative to proCoV2, which is a putative progenitor of SARS-CoV-2 inferred by Kumar *et al*. (2021) that differs from the widely used Wuhan-Hu-1 reference sequence by three mutations (C8782T, C18060T, and T28144C). Although requiring coverage of only ≥3 is relatively lenient, Table 1 shows that all sites with mutations have coverage ≥10. In addition, the mutations I called from the raw sequence data in Table 1 concord with those mentioned in Wang *et al*. (2020b). Again, this fact emphasizes that the information contained in the deleted sequencing data is largely present in Wang *et al*. (2020b), but because it was only published in a table in the chemistry journal *Small* rather than placed on the SRA, its existence was overlooked.

**Table 1.** Samples for which the SARS-CoV-2 sequence could be called at ≥ 90% of sites between 21,570 and 29,550, and the substitutions in this region relative to the putative SARS-CoV-2 progenitor proCoV2 inferred by Kumar *et al*. (2021). Numbers in parentheses after each substitution give the deep sequencing reads with each nucleotide identity.

| sample | fraction sites called (21570-29550) | patient group | substitutions relative to proCoV2 |
| --- | --- | --- | --- |
| A4 | 0.9266 | early outpatient | none |
| C1 | 0.9396 | early outpatient | G22081A (A=924, C=4, G=9), C28144T (C=6, T=1185), T29483G (C=1, G=45, T=1) |
| C2 | 0.9397 | early outpatient | C29095T (C=1, G=1, T=751) |
| C9 | 0.9005 | early outpatient | C28144T (C=3, T=823), G28514T (G=1, T=36) |
| D9 | 0.9051 | early outpatient | C28144T (C=4, T=1653) |
| D12 | 0.9400 | early outpatient | C28144T (C=8, T=2400) |
| E1 | 0.9223 | early outpatient | C28144T (T=125) |
| E5 | 0.9227 | early outpatient | C24034T (A=5, C=3, T=74), T26729C (C=12), G28077C (C=142, G=4) |
| E11 | 0.9321 | early outpatient | C25460T (C=2, T=246), C28144T (C=1, T=412) |
| F11 | 0.9054 | early outpatient | T25304A (A=9, T=1), C28144T (C=6, G=1, T=1328) |
| G1 | 0.9396 | early outpatient | none |
| G11 | 0.9112 | early outpatient | none |
| H9 | 0.9381 | early outpatient | C28144T (C=2, T=1254) |
| R11 | 0.9422 | hospital patient (Feb) | C21707T (T=401), C28144T (A=1, C=18, T=4265) |

I also determined the consensus sequence of the plasmid control used by Wang *et al*. (2020a) from the recovered sequencing data, and found that it had mutations C28144T and G28085T relative to proCoV2, which means that in the region of interest this control matches Wuhan-Hu-1 with the addition of G28085T. Since none of the viral samples in Table 1 contain G28085T and the samples that prove most relevant below also lack C28144T (which is a frequent natural mutation among early Wuhan sequences), plasmid contamination did not afflict the viral samples in the deleted sequencing project.

### Analysis of existing SARS-CoV-2 sequences emphasizes the perplexing discordance between collection date and distance to bat coronavirus relatives

To contextualize the viral sequences recovered from the deleted project, I first analyze early SARS-CoV-2 sequences already available in the GISAID database (Shu and McCauley 2017). The analyses described in this section are not entirely novel, but synthesize observations from multiple prior studies (Kumar *et al*. 2021; Pekar *et al.* 2021; Rambaut *et al.* 2020; Forster *et al.* 2020; Pipes *et al.* 2021) to provide key background.

Known human SARS-CoV-2 sequences are consistent with expansion from a single progenitor sequence (Kumar *et al.* 2021; Pekar *et al.* 2021; Rambaut *et al.* 2020; Forster *et al.* 2020; Pipes *et al.* 2021). However, attempts to infer this progenitor have been confounded by a perplexing fact: the earliest reported sequences from Wuhan are *not* the sequences most similar to SARS-CoV-2’s bat coronavirus relatives (Pipes *et al.* 2021). This fact is perplexing because although the proximal origin of SARS-CoV-2 remains unclear (i.e., zoonosis versus lab accident), all reasonable explanations agree that at a deeper level the SARS-CoV-2 genome is derived from bat coronaviruses (Lytras *et al*. 2021). One would therefore expect the first reported SARS-CoV-2 sequences to be the most similar to these bat coronavirus relatives—but this is not the case.

This conundrum is illustrated in Figure 2, which plots the collection date of SARS-CoV-2 sequences in GISAID versus the relative number of mutational differences from RaTG13 (Zhou *et al.* 2020b), which is the bat coronavirus with the highest full-genome sequence identity to SARS-CoV-2. The earliest SARS-CoV-2 sequences were collected in Wuhan in December, but these sequences are more distant from RaTG13 than sequences collected in January from other locations in China or even other countries (Figure 2). The discrepancy is especially pronounced for sequences from patients who had visited the Huanan Seafood Market (WHO 2021). All sequences associated with this market differ from RaTG13 by at least three more mutations than sequences subsequently collected at various other locations (Figure 2)—a fact that is difficult to reconcile with the idea that the market was the original location of spread of a bat coronavirus into humans. Importantly, all these observations also hold true if SARS-CoV-2 is compared to other related bat coronaviruses (Lytras *et al.* 2021) such as RpYN06 (Zhou *et al.* 2021) or RmYN02 (Zhou *et al.* 2020a) rather than RaTG13 (Figure S3).

**Figure 2.**
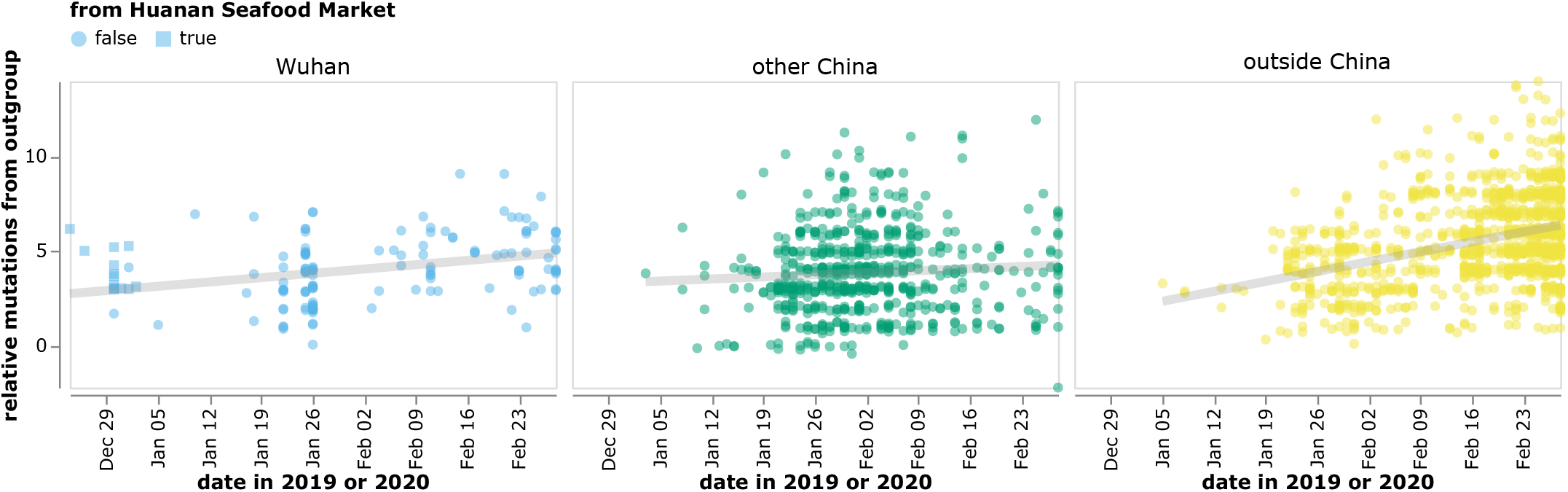
The reported collection dates of SARS-CoV-2 sequences in GISAID versus their relative mutational distances from the RaTG13 bat coronavirus outgroup. Mutational distances are relative to the putative progenitor proCoV2 inferred by Kumar *et al*. (2021). The plot shows sequences in GISAID collected no later than February 28, 2020. Sequences that the joint WHO-China report (WHO 2021) describes as being associated with the Wuhan Seafood Market are plotted with squares. Points are slightly jittered on the y-axis. Go to https://jbloom.github.io/SARS-CoV-2_PRJNA612766/deltadist.html for an interactive version of this plot that enables toggling of the outgroup to RpYN06 and RmYN02, mouseovers to see details for each point including strain name and mutations relative to proCoV2, and adjustment of the y-axis jittering. Static versions of the plot with RpYN06 and RmYN02 outgroups are in Figure S3.

This conundrum can be visualized in a phylogenetic context by rooting a tree of early SARS-CoV-2 sequences so that the progenitor sequence is closest to the bat coronavirus out-group. If we limit the analysis to sequences with at least two observations among strains collected no later than January 2020, there are three ways to root the tree in this fashion since there are three different sequences equally close to the outgroup (Figure 3, Figure S4). Importantly, none of these rootings place any Huanan Seafood Market viruses (or other Wuhan viruses from December 2019) in the progenitor node—and only one of the rootings has any virus from Wuhan in the progenitor node (in the leftmost tree in Figure 3, the progenitor node contains Wuhan/0126-C13/2020, which was reportedly collected on January 26, 2020). Therefore, inferences about the progenitor of SARS-CoV-2 based on comparison to related bat viruses are inconsistent with other evidence suggesting the progenitor is an early virus from Wuhan (Pipes *et al.* 2021).

**Figure 3.**
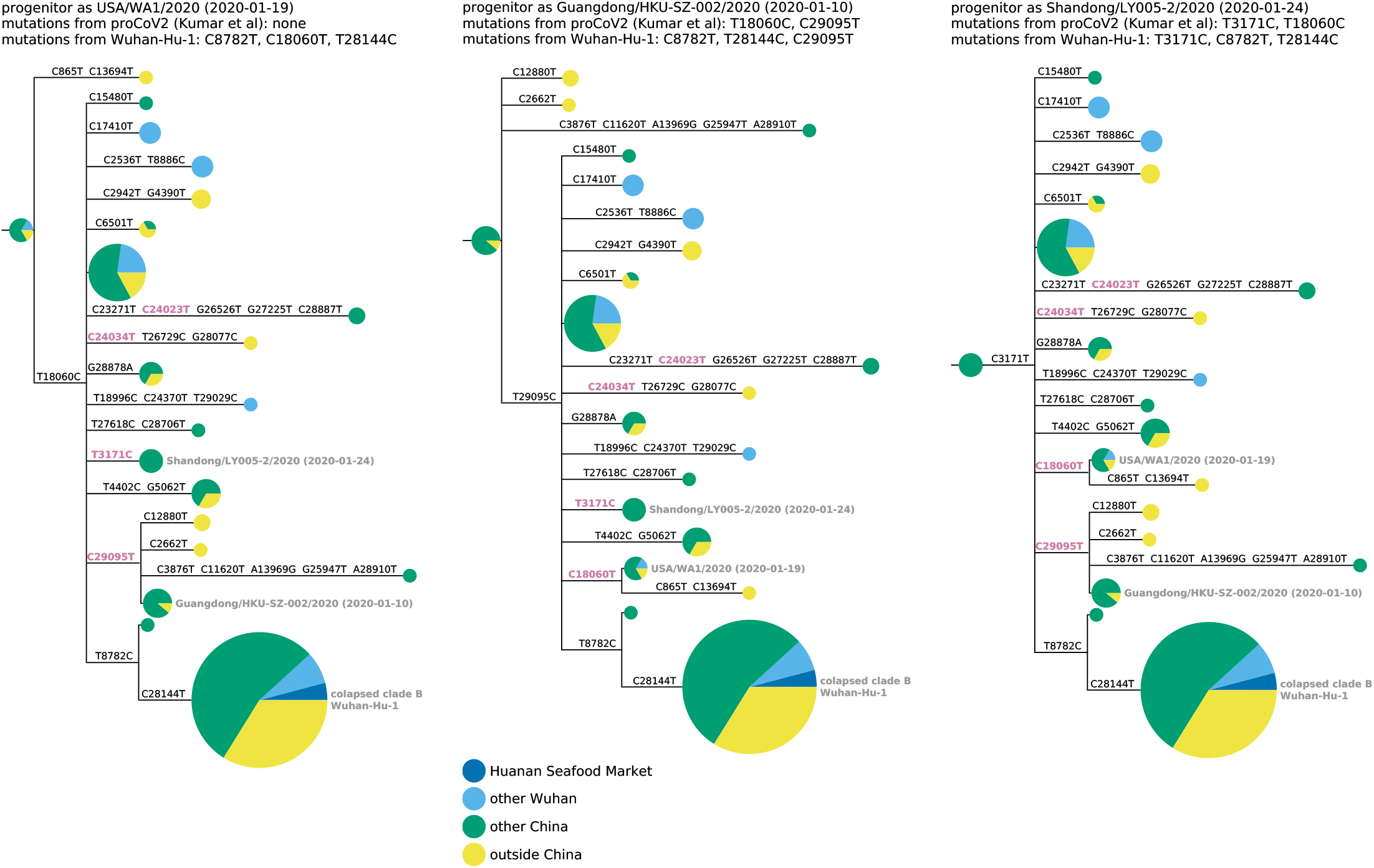
Phylogenetic trees of SARS-CoV-2 sequences in GISAID with multiple observations among viruses collected before Februrary, 2020. The trees are identical except they are rooted to make the progenitor each of the three sequences with highest identity to the RaTG13 bat coronavirus outgroup. Nodes are shown as pie charts with areas proportional to the number of observations of that sequence, and colored by where the viruses were collected. The mutations on each branch are labeled, with mutations towards the nucleotide identity in the outgroup in purple. The labels at the top of each tree give the first known virus identical to each putative progenitor, as well as mutations in that progenitor relative to proCoV2 (Kumar *et al.* 2021) and Wuhan-Hu-1. The monophyletic group containing C28144T is collapsed into a node labeled “clade B” in concordance with the naming scheme of Rambaut *et al*. (2020); this clade contains Wuhan-Hu-1. Figure S4 shows identical results are obtained if the outgroup is RpYN06 or RmYN02.

Several plausible explanations have been proposed for the discordance of phylogenetic rooting with evidence that Wuhan was the origin of the pandemic. Rambaut *et al*. (2020) suggest that viruses from the clade labeled “B” in Figure 3 may just “happen” to have been sequenced first, but that other SARS-CoV-2 sequences are really more ancestral as implied by phylogenetic rooting. Pipes *et al*. (2021) discuss the conundrum in detail, and suggest that phylogenetic rooting could be incorrect due to technical reasons such as high divergence of the outgroup or unusual mutational processes not captured in substitution models. Kumar *et al*. (2021) agree that phylogenetic rooting is problematic, and circumvent this problem by using an alternative algorithm to infer a progenitor for SARS-CoV-2 that they name proCoV2. Notably, proCoV2 turns out to be identical to one of the putative progenitors yielded by my approach in Figure 3 of simply placing the root at the nodes closest to the outgroup. However, neither the sophisticated algorithm of Kumar *et al*. (2021) nor my more simplistic approach explain *why* the progenitor should be so different from the earliest sequences reported from Wuhan.

Another explanation that I consider less plausible is offered by Garry (2021): that there were multiple zoonoses from distinct markets, with the Huanan Seafood Market being the source of viruses in clade B, and some other market being the source of viruses that lack T8782C and C28144T (Figure 3). However, this explanation requires positing zoonoses in two markets by two progenitors differing by just two mutations, which seems non-parsimonious in the absence of direct evidence for zoonosis in any market.

### Sequences recovered from the deleted project and better annotation of Wuhan-derived viruses help reconcile inferences about SARS-CoV-2’s progenitor

To examine if the sequences recovered from the deleted data set help resolve the conundrum described in the previous section, I repeated the analyses including those sequences. In the process, I noted another salient fact: four GISAID sequences collected in Guangdong that fall in a putative progenitor node are from two different clusters of patients who traveled to Wuhan in late December of 2019 and developed symptoms before or on the day that they returned to Guangdong, where their viruses were ultimately sequenced (Chan *et al.* 2020; Kang *et al.* 2020b). Since these patients were clearly infected in Wuhan even though they were sequenced in Guangdong, I annotated them separately from both the other Wuhan and other China sequences.

Repeating the analysis of the previous section with these changes shows that several sequences from the deleted project and all sequences from patients infected in Wuhan but sequenced in Guangdong are more similar to the bat coronavirus outgroup than sequences from the Huanan Seafood Market (Figure 4). This fact suggests that the market sequences, which are the primary focus of the genomic epidemiology in the joint WHO-China report (WHO 2021), are not representative of the viruses that were circulating in Wuhan in late December of 2019 and early January of 2020.

**Figure 4.**
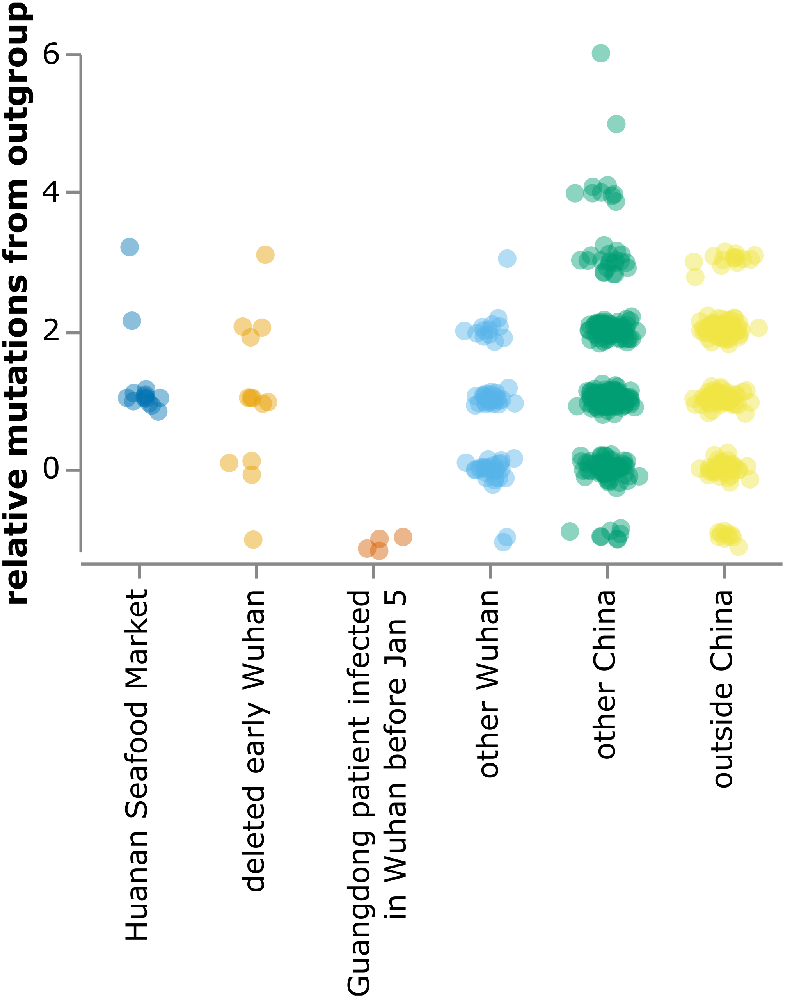
Relative mutational distance from RaTG13 bat coronavirus outgroup calculated *only* over the region of the SARS-CoV-2 genome covered by sequences from the deleted data set (21,570–29,550). The plot shows sequences in GISAID collected before February of 2020, as well as the 13 early Wuhan epidemic sequences in Table 1. Mutational distance is calculated relative to proCoV2, and points are jittered on the y-axis. Go to https://jbloom.github.io/SARS-CoV-2_PRJNA612766/deltadist_jitter.html for an interactive version of this plot that enables toggling the outgroup to RpYN06 or RmYN02, mouseovers to see details for each point, and adjustment of jittering.

Furthermore, it is immediately apparent that the discrepancy between outgroup rooting and the evidence that Wuhan was the origin of SARS-CoV-2 is alleviated by adding the deleted sequences and annotating Wuhan infections sequenced in Guangdong. The rooting of the middle tree in Figure 5 is now highly plausible, as half its progenitor node is derived from early Wuhan infections, which is more than any other equivalently large node. The first known sequence identical to this putative progenitor (Guangdong/HKU-SZ-002/2020) is from a patient who developed symptoms on January 4 while visiting Wuhan (Chan *et al.* 2020). This putative progenitor has three mutations towards the bat coronavirus outgroup relative to Wuhan-Hu-1 (C8782T, T28144C, and C29095T), and two mutations relative to proCoV2 (T18060C away from the outgroup and C29095T towards the outgroup). The leftmost tree in Figure 5, which has a progenitor identical to proCoV2 (Kumar *et al.* 2021) also looks plausible, with some weight from Wuhan sequences. However, analysis of this rooting is limited by the fact that the defining C18060T mutation is in a region not covered in the deleted sequences. The rightmost tree in Figure 5 looks less plausible, as it has almost no weight from Wuhan and the first sequence identical to its progenitor was not collected until January 24.

**Figure 5.**
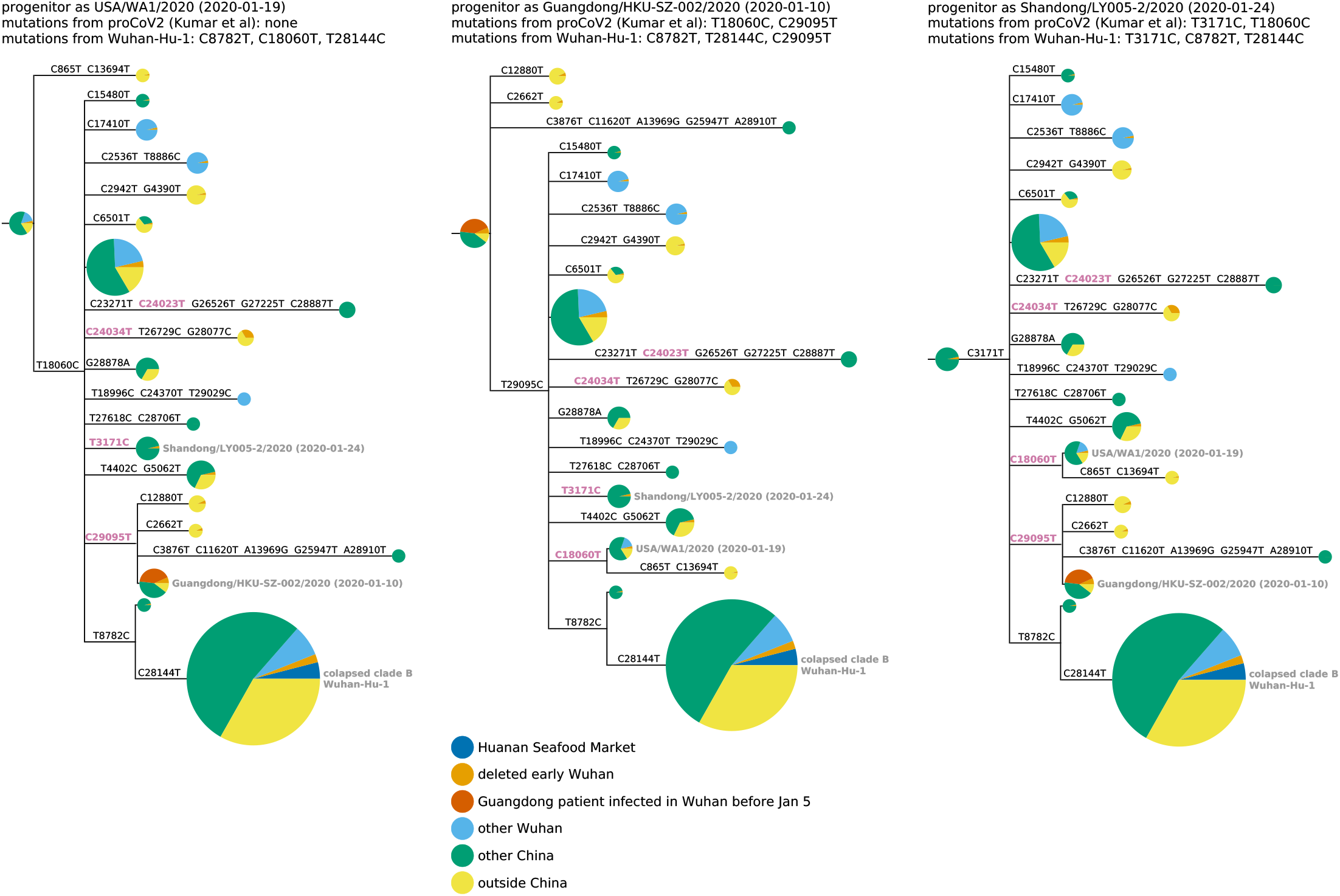
Phylogenetic trees like those in Figure 3 with the addition of the early Wuhan epidemic sequences from the deleted data set, and Guangdong patients infected in Wuhan prior to January 5 annotated separately. Because the deleted sequences are partial, they cannot all be placed unambiguously on the tree. Therefore, they are added to each compatible node proportional to the number of sequences already in that node. The deleted sequences with C28144T (clade B) or C29095T (putative progenitor in middle tree) can be placed relatively unambiguously as defining mutations occur in the sequenced region, but those that lack either of these mutations are compatible with a large number of nodes including the proCoV2 putative progenitor. Figure S4 demonstrates that the results are identical if RpYN06 or RmYN02 is instead used as the outgroup.

We can also qualitatively examine the three progenitor placements in Figure 5 using the principle employed by Worobey *et al*. (2020) to help evaluate scenarios for emergence of SARS-CoV-2 in Europe and North America: namely that during a growing outbreak, a progenitor is likely to give rise to multiple branching lineages. This principle is especially likely to hold for the scenarios in Figure 5, since there are multiple individuals infected with each putative progenitor sequence, implying multiple opportunities to transmit descendants with new mutations. Using this qualitative principle, the middle scenario in Figure 5 seems most plausible, the leftmost (proCoV2) scenario also seems plausible, and the rightmost scenario seems less plausible. I acknowledge these arguments are purely qualitative and lack the formal statistical analysis of Worobey *et al*. (2020)—but as discussed below, there may be wisdom in qualitative reasoning when there are valid concerns about the nature of the underlying data.

## Discussion

I have identified and recovered a deleted set of partial SARS-CoV-2 sequences from the early Wuhan epidemic. Analysis of these sequences leads to several conclusions. First, they provide further evidence Huanan Seafood Market sequences that were the focus of the joint WHO-China report (WHO 2021) are not representative of all SARS-CoV-2 in Wuhan early in the epidemic. The deleted data as well as existing sequences from Wuhan-infected patients hospitalized in Guangdong show early Wuhan sequences often carried the T29095C mutation and were less likely to carry T8782C / C28144T than sequences in the joint WHO-China report (WHO 2021). Second, given current data, there are two plausible identities for the progenitor of all known SARS-CoV-2. One is proCoV2 described by Kumar *et al*. (2021), and the other is a sequence that carries three mutations (C8782T, T28144C, and C29095T) relative to Wuhan-Hu-1. Crucially, both putative progenitors are three mutations closer to SARS-CoV-2’s bat coronavirus relatives than sequences from the Huanan Seafood Market. Note also that the progenitor of all known SARS-CoV-2 sequences could still be downstream of the sequence that infected patient zero—and it is possible that the future discovery of additional early SARS-CoV-2 sequences could lead to further revisions of inferences about the earliest viruses in the outbreak.

The fact that this informative data set was deleted suggests implications beyond those gleaned directly from the recovered sequences. Samples from early outpatients in Wuhan are a gold mine for anyone seeking to understand spread of the virus. Even my analysis of 13 partial sequences is revealing, and it clearly would have been more scientifically informative to fully sequence all 34 samples rather than delete the partial sequence data. There is no obvious scientific reason for the deletion: the sequences are concordant with the samples described in Wang *et al*. (2020a,b), there are no corrections to the paper, the paper states human subjects approval was obtained, and the sequencing shows no evidence of plasmid or sample-to-sample contamination. After I e-mailed the NIH the original version of this manuscript, they sent me the e-mails requesting deletion of the data, which are in Figure 6. While I cannot rule out that the authors posted the data on some unknown website, the sequences are not in GISAID, NCBI, or any database used by the joint WHO-China report. Therefore, even though the sequencing data were on the Google Cloud (as described above) and the mutations were listed in a table in the *Small* paper by Wang *et al*. (2020b), the practical consequence of removing the data from the SRA was that nobody was aware these sequences existed. Particularly in light of the directive that labs destroy early samples (Pingui 2020) and multiple orders requiring approval of publications on COVID-19 (China CDC 2020; Kang *et al.* 2020a), this suggests a less than wholehearted effort to maximize information about viral sequences from early in the Wuhan epidemic.

**Figure 6.**
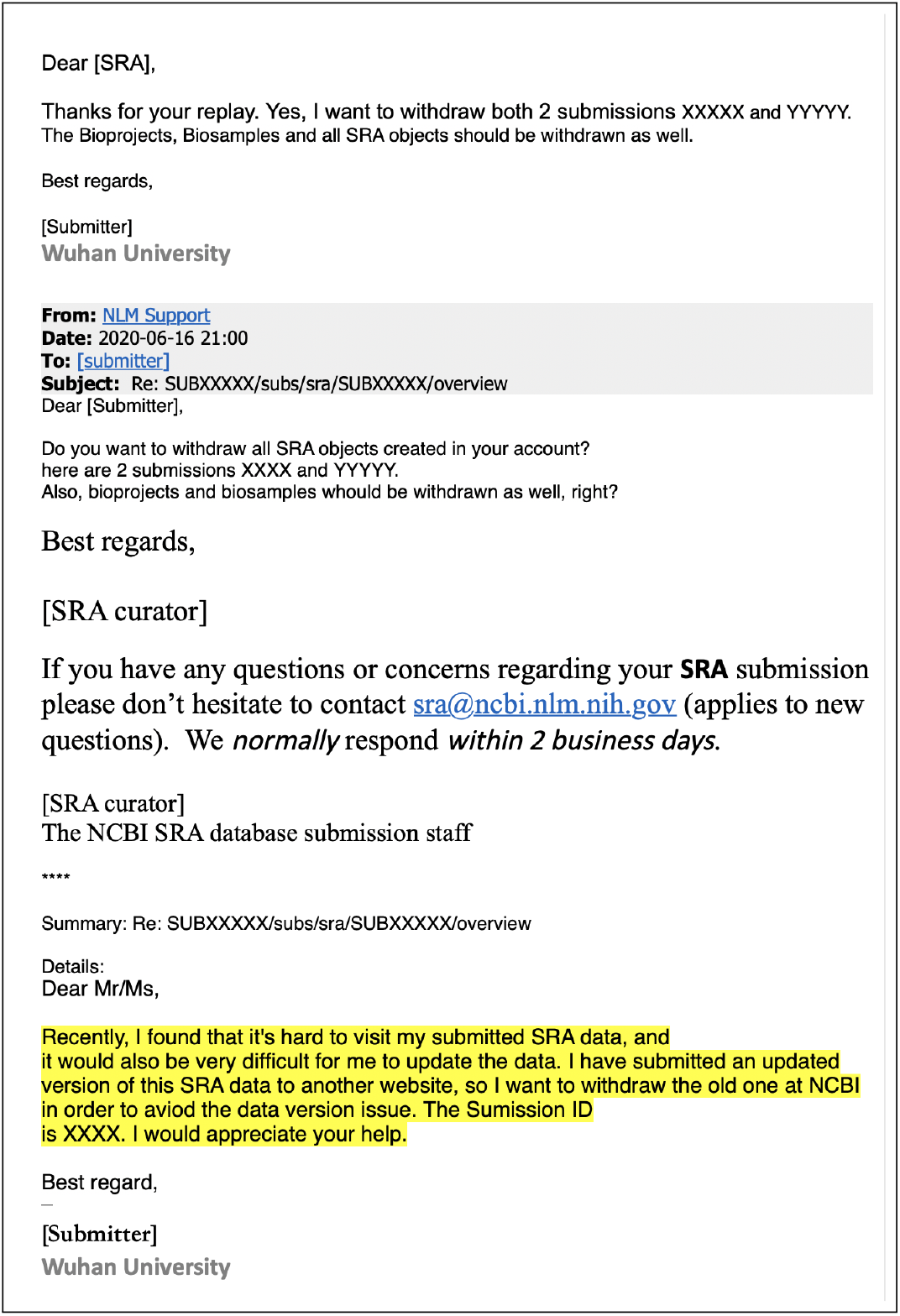
A redacted version of the e-mails from Wuhan University to the SRA staff requesting deletion of the sequencing data. This e-mail was provided to me by the NIH’s NCBI Director Stephen Sherry on June, 19, 2021, the day after I emailed the NIH an advance copy of this manuscript. The redactions and highlighting were done by the NIH, and I am showing the e-mail exactly as it was provided to me.

Another important implication is that genomic epidemiology studies of early SARS-CoV-2 need to pay as much attention to the provenance and annotation of the underlying sequences as technical considerations. There has been substantial scientific effort expended on topics such as phylogenetic rooting (Pipes *et al*. 2021; Morel *et al*. 2021), novel algorithms (Kumar *et al*. 2021), and correction of sequencing errors (Turakhia *et al.* 2020). Future studies should devote equal effort to going beyond the annotations in GISAID to carefully trace the location of patient infection and sample sequencing. The potential importance of such work is revealed by the observation that many of the sequences closest to SARS-CoV-2’s bat coronavirus relatives are from early patients who were infected in Wuhan, but then sequenced in and attributed to Guangdong.

There are several caveats to this study. Most obviously, the sequences recovered from the deleted data set are partial and lack full metadata. Therefore, it is impossible to unambiguously place them phylogenetically, or determine exactly when they were collected. However, little can be done to mitigate this caveat beyond my unsuccessful attempt to contact the corresponding authors of Wang *et al*. (2020a). It is also important to note that my phylogenetic analyses use relatively simple methods to draw qualitative conclusions without formal statistical testing. Further application of more advanced methods would be a welcome advance. However, qualitative and visual analyses do have advantages when the key questions relate more to the underlying data than the sophistication of the inferences. Finally, both plausible putative progenitors require that an early mutation to SARS-CoV-2 was a reversion towards the bat coronavirus outgroups (either C18060T or C29095T) on a branch that subsequently gave rise to multiple distinct descendants. Such a scenario can only be avoided by invoking recombination very early in the pandemic, which is not entirely implausible for a coronavirus (Boni *et al.* 2020). However, because the outgroups have ∼ 4% nucleotide divergence from SARS-CoV-2, a mutation towards the outgroup is also entirely possible. Of course, future identification of additional early sequences could fully resolve these questions.

More broadly, the approach taken here suggests it may be possible to learn more about the origin or early spread of SARS-CoV-2 even without an international investigation. I suggest it could be worthwhile for the NIH to review e-mail records to identify other SRA deletions. Importantly, SRA deletions do not imply any malfeasance: there are legitimate reasons for removing sequencing runs, and the SRA houses >13-million runs making it infeasible for its staff to validate the rationale for all requests. However, the current study suggests that at least in one case, the trusting structures of science permitted a data deletion that obscured sequences relevant to the early spread of SARS-CoV-2 in Wuhan. A careful re-evaluation of other archived forms of scientific communication, reporting, and data could shed light on additional overlooked information relevant to the early emergence of the virus.

## Methods

### Code and data availability

The computer code and input data necessary to reproduce all analyses described in this paper are available on GitHub at https://github.com/jbloom/SARS-CoV-2_PRJNA612766. This GitHub repository includes a `Snakemake` (Mölder *et al.* 2021) pipeline that fully automates all steps in the analysis except for downloading of sequences from GISAID, which must be done manually as described in the GitHub repository’s README in order to comply with GISAID data sharing terms.

The deleted SRA files recovered from the Google Cloud are all available at https://github.com/jbloom/SARS-CoV-2_PRJNA612766/tree/main/results/sra_downloads. I have suffixed the file extension `.sra` to all these files. The consensus sequences recovered from these deleted SRA files are linked to in the relevant Methods subsection below.

### Archiving of key weblinks

I have digitally archived key weblinks in the Wayback Machine, including a subset of the SRA files from PRJNA612766 on the Google Cloud:

- The first supplementary table of Farkas *et al*. (2020) is archived at https://web.archive.org/web/20210502130356/ https://dfzljdn9uc3pi.cloudfront.net/2020/9255/1/Supplementary_Table_1.xlsx.
- SRR11313485: https://storage.googleapis.com/nih-sequence-read-archive/run/SRR11313485/SRR11313485
- SRR11313486: https://storage.googleapis.com/nih-sequence-read-archive/run/SRR11313486/SRR11313486
- SRR11313274: https://storage.googleapis.com/nih-sequence-read-archive/run/SRR11313274/SRR11313274
- SRR11313275: https://storage.googleapis.com/nih-sequence-read-archive/run/SRR11313275/SRR11313275
- SRR11313285: https://storage.googleapis.com/nih-sequence-read-archive/run/SRR11313285/SRR11313285
- SRR11313286: https://storage.googleapis.com/nih-sequence-read-archive/run/SRR11313286/SRR11313286
- SRR11313448: https://storage.googleapis.com/nih-sequence-read-archive/run/SRR11313448/SRR11313448
- SRR11313449: https://storage.googleapis.com/nih-sequence-read-archive/run/SRR11313449/SRR11313449
- SRR11313427: https://storage.googleapis.com/nih-sequence-read-archive/run/SRR11313427/SRR11313427
- SRR11313429: https://storage.googleapis.com/nih-sequence-read-archive/run/SRR11313429/SRR11313429

### Recovery of SRA files from deleted project PRJNA612766

I parsed the first supplementary table of Farkas *et al*. (2020) to extract the accessions for sequencing runs for deleted SRA BioProject PRJNA612766. By cross-referencing the samples described in this table to Wang *et al*. (2020a,b), I identified the accessions corresponding to the 34 early outpatient samples who were positive, as well as the accessions corresponding to the 16 hospitalized patient samples from February. Samples had both 10 minute and 4 hour sequencing runtime accessions, which were combined in the subsequent analysis. I also identified the samples corresponding to the high-copy plasmid controls to enable analysis of the plasmid sequence to rule out contamination. The code used to parse the Excel table is available as a Jupyter notebook at https://github.com/jbloom/SARS-CoV-2_PRJNA612766/tree/main/manual_analyses/PRJNA612766. I recovered the SRA files from the Google Cloud by using `wget` to download files with from paths like https://storage.googleapis

.com/nih-sequence-read-archive/run/SRR11313485/SRR11313485. Note that I cannot guarantee that these Google Cloud links will remain active, as my analyses of other deleted SRA runs (beyond the scope of this study) indicates that only sometimes are deleted SRA files still available via the Google Cloud. For this reason, key runs have been archived on the Wayback Machine as described above, and all downloaded SRA files relevant to this study are included in the GitHub repository. Note also that as described in this paper’s main text, two SRA files could not be downloaded from the Google Cloud using the aforementioned method, and so are not part of this study.

### FASTQ files for SRR11313490 and SRR11313499

I was not able to download SRA files for two runs, SRR11313490 and SRR11313499, from the Google Cloud. After I posted the initial version of this pre-print, I was contacted by three different individuals who had uncovered FASTQ or FASTA files for these two missing runs that were downloaded from the SRA prior to the deletion of PRJNA612766. For revised versions of the manuscript, I included analysis of the data for these two runs that I obtained from the links provided at https://github.com/lifebit-ai/SARS-CoV-2/blob/master/assets/ucsc/aws_https_links.txt. Inclusion of data for these two runs did not appreciably change the results of the analysis relative to that in the original version of the pre-print, since both runs corresponded to low coverage samples for which meaningful viral genetic information could not be obtained.

### Alignment of recovered reads and calling of consensus sequences

The downloaded SRA files were converted to FASTQ files using `fasterq-dump` from the SRA Toolkit. The FASTQ files were pre-processed with `fastp` (Chen *et al.* 2018) to trim reads and remove low-quality ones (the exact settings using in this pre-processing are specified in the `Snakemake` file in the GitHub repository).

The reads in these FASTQ files were then aligned to a SARS-CoV-2 reference genome using `minimap2` (Li 2018) with default settings. The reference genome used for the entirety of this study is proCoV2 (Kumar *et al.* 2021), which was generated by making the following three single-nucleotide changes to the Wuhan-Hu-1 reference (ASM985889v2) available on NCBI: C8782T, C18060T, and T28144C.

I processed the resulting alignments with `samtools` and `pysam` to determine the coverage at each site by aligned nucleotides with a quality score of at least 20. I also masked (set to zero coverage) all sites overlapped by the primers used by Wang *et al*. (2020a) to PCR the amplicons used for sequencing. These coverage plots are in Figure S1 and Figure S2; the legends of these figures also link to interactive versions of the plots that enable zooming and mouseovers to get statistics for specific sites. I called the consensus sequence at a site if this coverage was ≥ 3 and >80% of the reads agreed on the identity. These consensus sequences over the entire SARS-CoV-2 genome are available at https://github.com/jbloom/SARS-CoV-2_PRJNA612766/raw/main/results/consensus/consensus_seqs.csv; note that they are mostly `N` nucleotides since the sequencing approach of Wang *et al*. (2020a) only covers part of the genome.

I only used the recovered consensus sequences in the downstream analyses if it was possible to call the consensus identity at ≥ 90% of the sites in the region of interest from site 21,570 to 29,550. These are the sequences listed in Table 1, and as described in that table, all mutation calls were at sites with coverage ≥ 10. These sequences in the region of interest (21,570 to 29,550) are available at https://github.com/jbloom/SARS-CoV-2_PRJNA612766/blob/main/results/recovered_seqs.fa.

### Bat coronavirus outgroup sequences

For analyses that involved comparisons to SARS-CoV-2’s bat coronavirus relatives (Lytras *et al.* 2021), the bat coronavirus sequences were manually downloaded from GISAID (Shu and McCauley 2017). The sequences used were RaTG13 (Zhou *et al.* 2020b), RmYN02 (Zhou *et al*. 2020a), and RpYN06 (Zhou *et al.* 2021)—although the multiple sequence alignment of these viruses to SARS-CoV-2 also contains PrC31 (Li *et al*. 2021), which was not used in the final analyses as it more diverged from SARS-CoV-2 than the other three bat coronaviruses at a whole-genome level. The GISAID accessions for these sequences are listed at https://github.com/jbloom/SARS-CoV-2_PRJNA612766/blob/main/data/comparator_genomes_gisaid/accessions.txt, and a table acknowledging the labs and authors is at https://github.com/jbloom/SARS-CoV-2_PRJNA612766/blob/main/data/comparator_genomes_gisaid/acknowledgments.csv. Sites in SARS-CoV-2 were mapped to their corresponding nucleotide identities in the bat coronavirus outgroups via a multiple sequence alignment of proCoV2 to the bat coronaviruses generated using `mafft` (Katoh and Standley 2013).

### Curation and analysis of early SARS-CoV-2 sequences from GISAID

For the broader analyses of existing SARS-CoV-2 sequences, I downloaded all sequences from collected prior to March of 2020 from GISAID. The accessions of these sequences are at https://github.com/jbloom/SARS-CoV-2_PRJNA612766/blob/main/data/gisaid_sequences_through_Feb2020/accessions.txt, and a table acknowledging the labs and authors is at https://github.com/jbloom/SARS-CoV-2_PRJNA612766/blob/main/data/gisaid_sequences_through_Feb2020/acknowledgments.csv.

I then used `mafft` (Katoh and Standley 2013) to align these sequences to the proCoV2 reference described above, stripped any sites that were gapped relative to the reference, and filtered the sequences using the following criteria:

- I removed any sequences collected after February 28, 2020.
- I removed any sequences that had ≥ 4 mutations within any 10-nucleotide stretch, as such runs of mutations often indicate sequencing errors.
- I removed any sequence for which the alignment covered < 90% of the proCoV2 sequence.
- I removed any sequence with ≥ 15 mutations relative to the reference.
- I removed any sequence with ≥5,000 ambiguous nucleotides.

I then annotated the sequences using some additional information. First, I annotated sequences based on the joint WHO-China report (WHO 2021) and also Zhu *et al*. (2020) to keep only one representative from multiply sequenced patients, and to indicate which sequences were from patients associated with the Huanan Seafood Market. My version of these annotations is at https://github.com/jbloom/SARS-CoV-2_PRJNA612766/blob/main/data/WHO_China_Report_Dec2019_cases.yaml. Next, I identified some sequences in the set that were clearly duplicates from the same patient, and removed these. The annotations used to remove these duplicates are at https://github.com/jbloom/SARS-CoV-2_PRJNA612766/blob/main/data/seqs_to_exclude.yaml. Finally, I used information from Chan *et al*. (2020) and Kang *et al*. (2020b) to identify patients who were infected in Wuhan before January 5 of 2020, but ultimately sequenced in Guangdong: these annotations are at https://github.com/jbloom/SARS-CoV-2_PRJNA612766/blob/main/data/Wuhan_exports.yaml.

I next removed any of the handful of mutations noted by Turakhia *et al*. (2020) to be lab artifacts that commonly afflict SARS-CoV-2 sequences. I also limited the analyses to the region of the genome that spans from the start of the first coding region (ORF1ab) to the end of the last (ORF10), because I noticed that some sequences had suspicious patterns (such as many mutations or runs of mutations) near the termini of the genome.

The plot in Figure 2 contains all of the GISAID sequences after this filtering. The plot in Figure 4 shows the filtered GISAID sequences collected before February of 2020 plus the 13 good coverage recovered partial early outpatient sequences (Table 1), considering only the region covered by the partial sequences (21,570 to 29,550).

### Phylogenetic analyses

The phylogenetic trees were inferred using the GISAD sequences after the filtering and annotations described above, only considering sequences with ≥ 95% coverage over the region of interest that were collected before February of 2020. In addition, after generating this sequence set I removed any sequence variants with a combination of mutations that was not observed at least twice so the analysis only includes multiply observed sequence variants. A file indicating the unique sequences used for the phylogenetic analysis, their mutations relative to proCoV2, and other sequences in that cluster is at https://github.com/jbloom/SARS-CoV-2_PRJNA612766/blob/main/results/phylogenetics/all_alignment.csv.

I then used `IQ-Tree` (Minh *et al.* 2020) to infer a maximum-likelihood phylogenetic tree using a GTR nucleotide substitution model with empirical nucleotide frequencies, and collapsing zero-length branches to potentially allow a multifurcating tree. The inference yielded the tree topology and branch lengths shown in all figures in this study with phylogenetic trees. I then rendered the images of the tree using `ETE 3` (Huerta-Cepas *et al.* 2016), manually re-rooting the tree to place the first (progenitor) node at each of the three nodes that have the highest identity to the bat coronavirus outgroup. In these images, node sizes are proportional to the number of sequences in that node, and are colored in proportion to the location from which those sequences are derived. As indicated in the legend to Figure 3, the node containing the monophyletic set of sequences with C28144T is collapsed into a single node in the tree images.

For the trees in which I added the recovered sequences from the deleted data set (Figure 5), the actual trees are exactly the same as those inferred using the GISAID sequences above. The difference is that the sequences from the deleted data set are then added to each node with which they are compatible given their mutations in an amount proportional to the size of the node, the logic being that a sequence is more likely to fall into larger clusters.

### Interactive versions of some figures

Interactive versions of some figures are available at https://jbloom.github.io/SARS-CoV-2_PRJNA612766/, and were created using `Altair` (Van-derPlas *et al.* 2018)

## Acknowledgments

I thank the citizens and scientists on Twitter who helped inspire this study and inform its background through their analyses and discussions of the early spread of SARS-CoV-2. I thank Brendan Larsen for pointing out that I failed to mask the primer binding site regions in the analysis described in the original pre-print (this has been fixed in this version). I thank Stephen Goldstein and anonymous commenter for providing critiques of the original version of the pre-print on *bioRxiv* that have helped me revise the manuscript. I thank the scientists and labs who contributed sequences used in this study to the GISAID database; the names of these scientists and labs are listed in the tables linked to in the Methods. The scientific computing infrastructure used in this work was supported by the NIH Office of Research Infrastructure Programs under S10OD028685. The author is an Investigator of the Howard Hughes Medical Institute.

