## Supplementary Material for "Recovery of deleted deep sequencing data sheds more light on the early Wuhan SARS-CoV-2 epidemic"

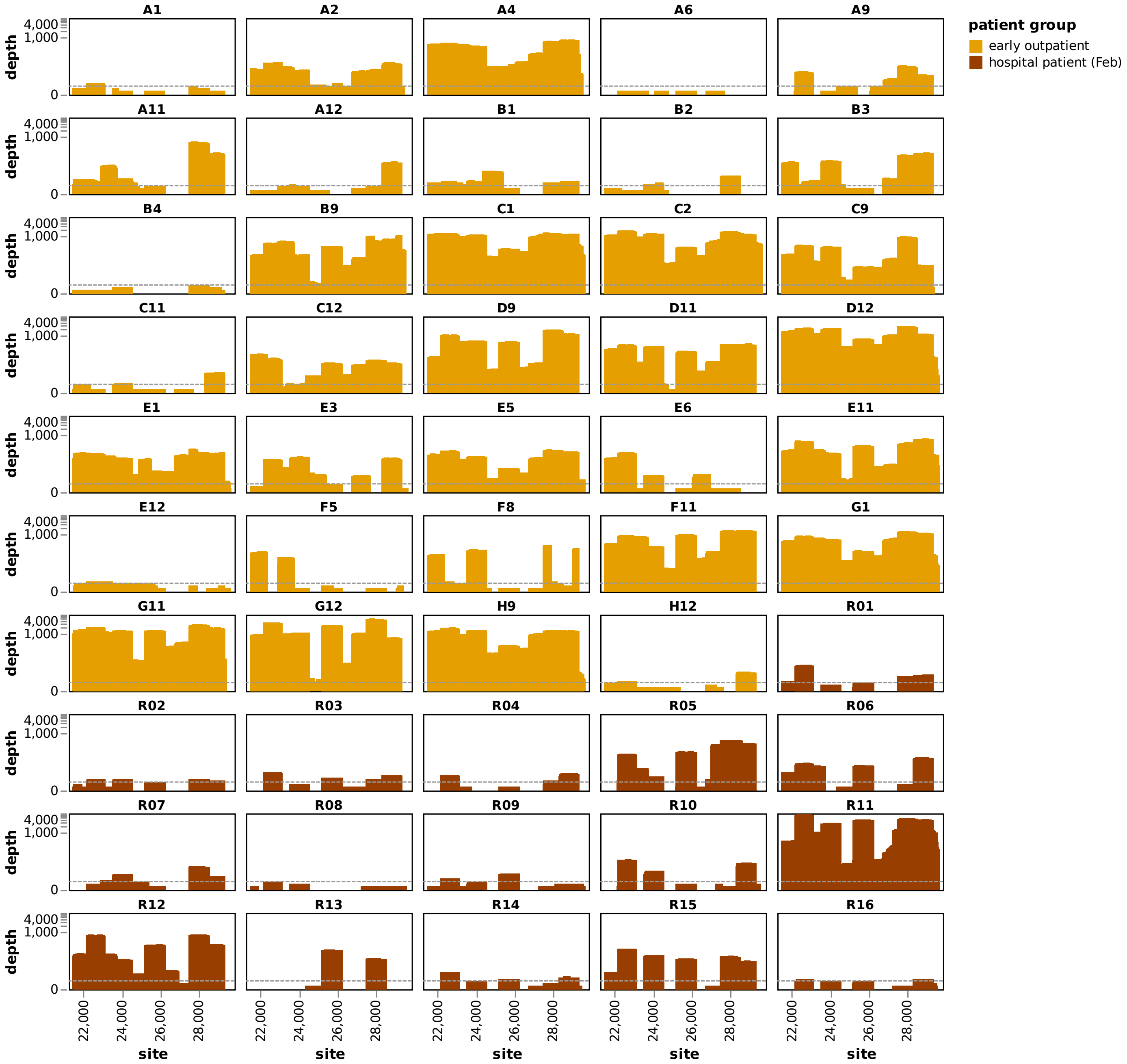


**Figure S1** Sequencing depth over the SARS-CoV-2 genome from site 21,570 to 29,550 for the 34 virus-positive early epidemic samples and the 16 samples from hospitalized patients in February. Depth is the number of aligned reads that cover that site with a quality score ≥20. The dashed gray line is the minimum coverage required to call a consensus identity at a site. Note that the y-axis uses a symlog scale. An interactive version of this plot that enables zooming into specific site ranges and mouseovers to see read count statistics at each site is at https://jbloom.github.io/SARS-CoV-2_PRJNA612766/coverage_region.html. A version of the plot where the x-axis spans the entire SARS-CoV-2 genome is in Figure S2.

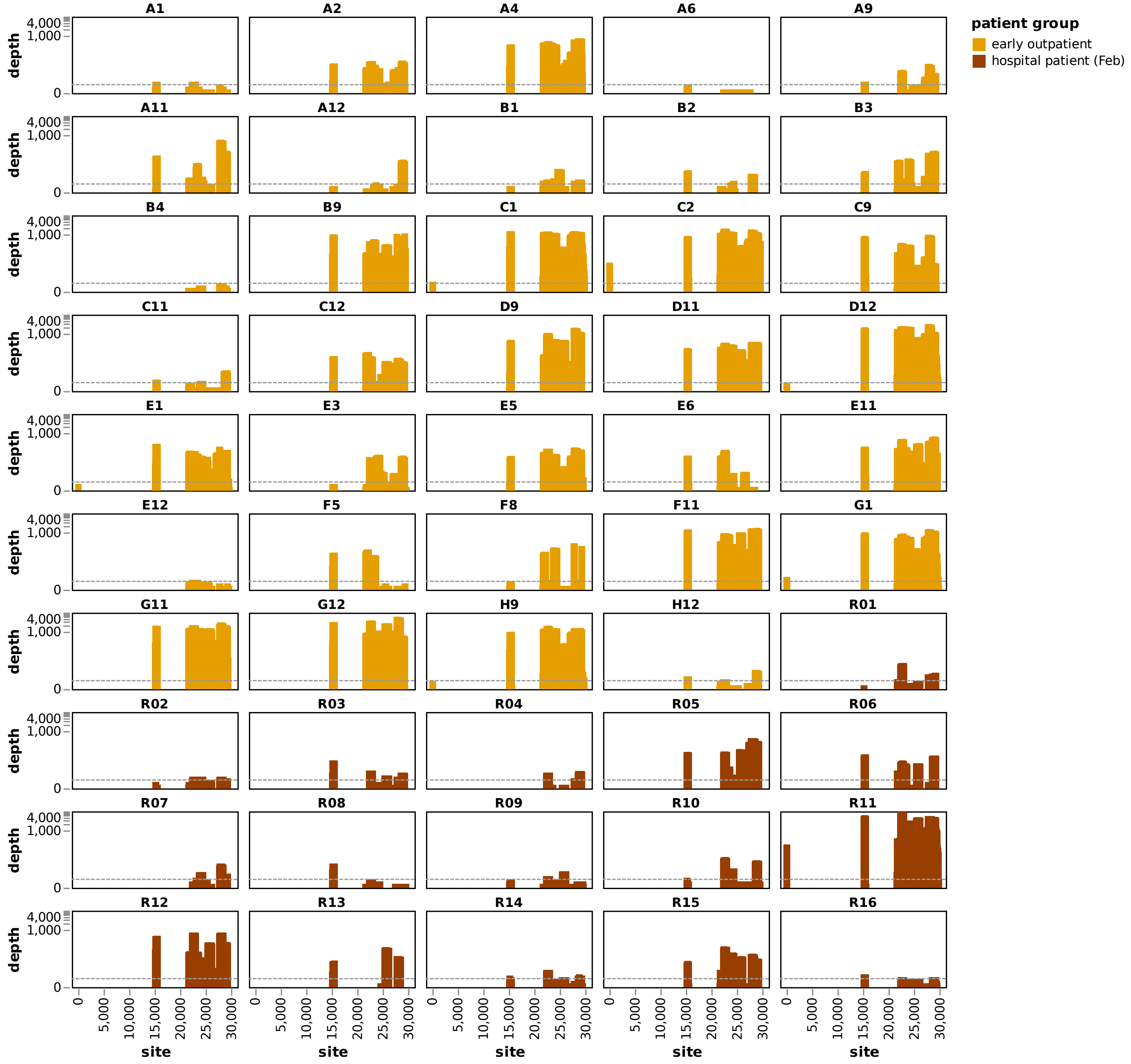


**Figure S2** A version of Figure S1 that shows coverage over the full length of the SARS-CoV-2 genome. An interactive version of this plot is at https://jbloom.github.io/SARS-CoV-2_PRJNA612766/coverage_all.html.

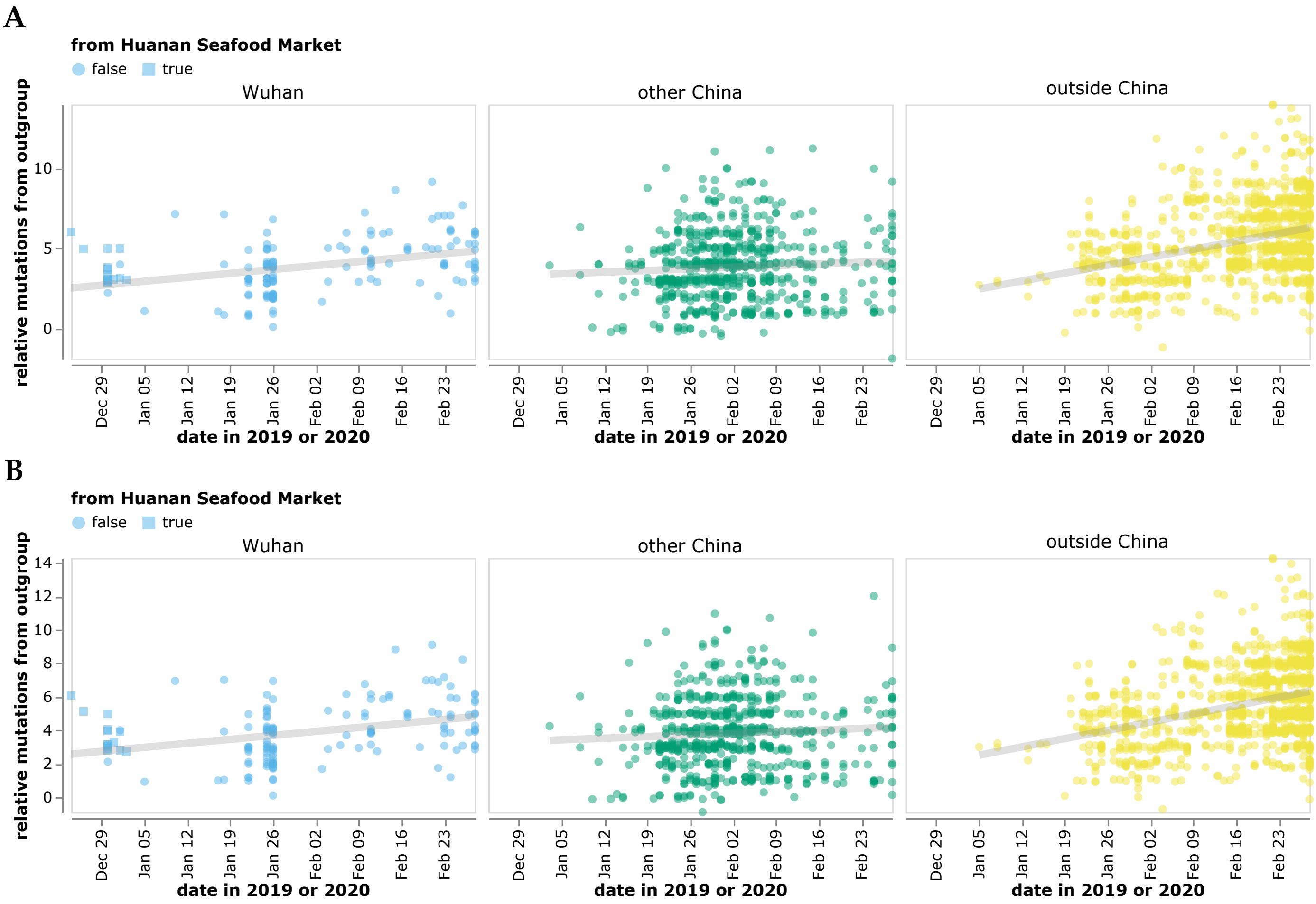


**Figure S3** Versions of Figure 2 but calculating the relative mutational distances using an outgroup of (A) RpYN06 or (B) RmYN02.

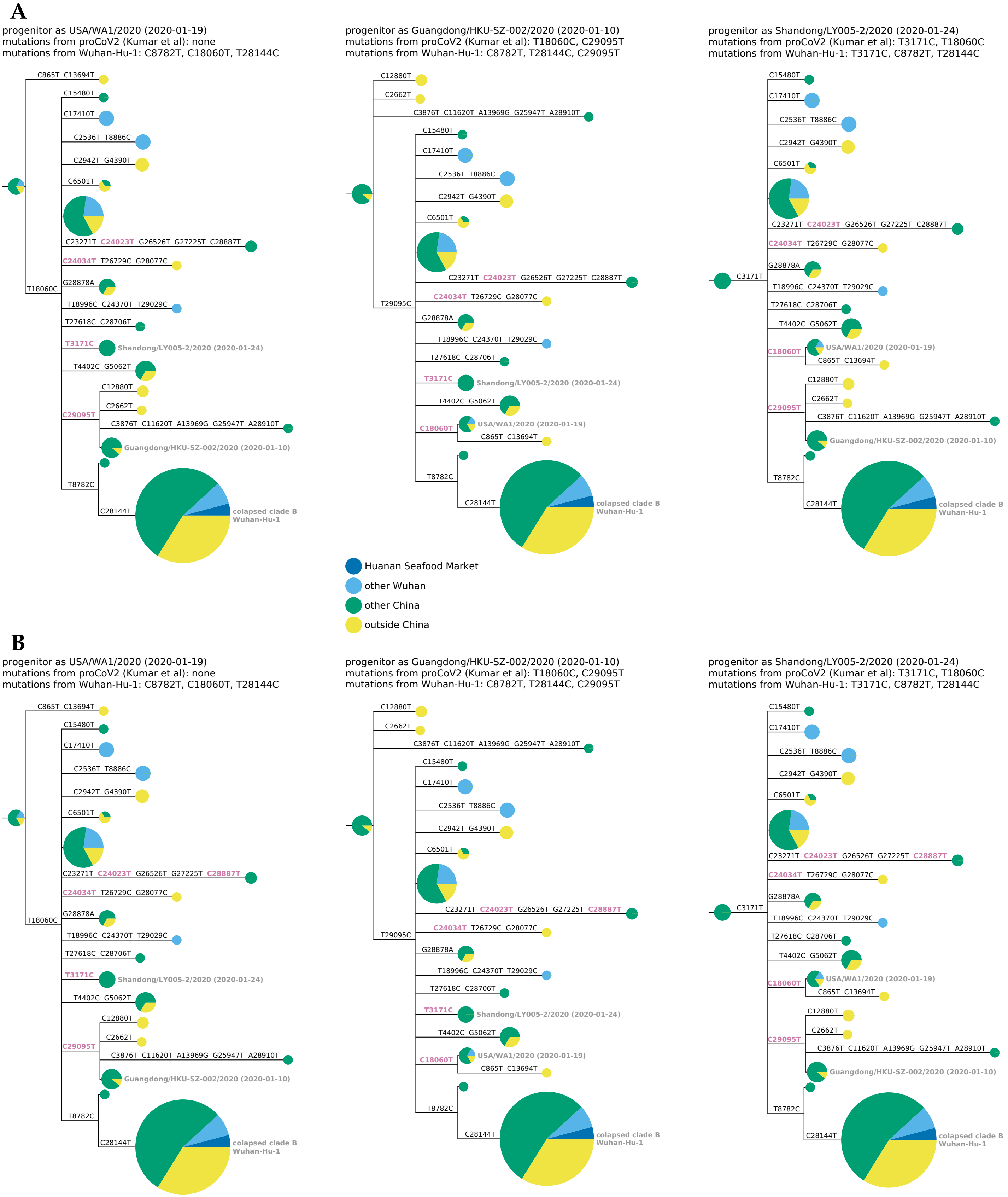


**Figure S4** A version of Figure 3 but rooting using an outgroup of (A) RpYN06 or (B) RmYN02 outgroups. The tree topologies are identical to those obtained using RaTG13 in Figure 3, with the only differences being a few minor changes in which mutations on branches are towards the outgroup (purple mutation labels).
